# New barcoded primers for efficient retrieval of cercozoan sequences in high-throughput environmental diversity surveys, with emphasis on worldwide biological soil crusts

**DOI:** 10.1101/171611

**Authors:** Anna Maria Fiore-Donno, Christian Rixen, Martin Rippin, Karin Glaser, Elena Samolov, Ulf Karsten, Burkhard Becker, Michael Bonkowski

## Abstract

We describe the performance of a new metabarcoding approach to investigate the environmental diversity of a prominent group of widespread unicellular organisms, the Cercozoa. Cercozoa is an immensely large group of protists and although it may dominate in soil and aquatic ecosystems, its environmental diversity remains undersampled. We designed PCR primers targeting the hyper-variable region V4 of the small subunit ribosomal RNA (SSU or 18S) gene, which is the recommended barcode marker for Cercozoa. The length of the amplified fragment (ca. 350 bp) is suitable for Illumina MiSeq, the most cost-effective platform for molecular environmental surveys. We provide barcoded primers, an economical alternative to multiple libraries for multiplex sequencing of over a hundred samples. *In silico*, our primers matched 68% of the cercozoan sequences of the reference database and performed better than previously proposed new generation sequencing primers. In mountain grasslands soils and in biological soil crusts from a variety of climatic regions, we were able to detect cercozoan sequences encompassing nearly the whole range of the phylum. We obtained 901 OTUs at 97% similarity threshold from 26 samples, with ca. 50,000 sequences per site, and only 8% of non-cercozoan sequences. We could contribute to a further increase of the diversity of Cercozoa, since only 43% of the OTUs were 97-100% similar to any known sequence. Our study thus provides an advanced tool for cercozoan metabarcoding and to investigate their diversity and distribution in the environment.

## 1. Introduction

Environmental molecular surveys are becoming the state-of-the-art tool to conduct worldwide inventories of microorganisms, often revealing a hidden and unsuspected diversity (de Vargas *et al.* 2015). A step forward in microbial ecology has been made due to the advent of high-throughput sequencing and especially Illumina sequencing: by providing more than ten millions sequences per run, the depth needed to conduct large-scale biodiversity surveys can be achieved. Despite this, eukaryotic diversity is still understudied compared to that of prokaryotes. Protists in general remain undersampled, partly because they are a heterogeneous assemblage of unrelated lineages, spanning an immense genetic diversity (Pawlowski *et al.* 2012).

The method of choice for protistan surveys and barcoding consists in amplifying fragments of the small subunit ribosomal RNA (SSU or 18S) gene, the gene with the largest reference database (Pawlowski *et al.* 2012). The V4 hypervariable region located at the end of the first third of the SSU has been recommended as the universal protistan barcode (Pawlowski *et al.* 2012), as useful as the full-length gene for assessing protistan diversity (Hu *et al.* 2015). PCR-free approaches, such as direct sequencing of whole DNAs or cDNAs, at present result in many unrecognizable genes because free-living species of protists are underrepresented in the reference molecular databases, which are typically dominated by fungi, plants and animals (del Campo *et al.* 2014).

So-called “universal eukaryotic primers” have been designed to target a wide range of eukaryotes (Bates *et al.* 2013; Massana *et al.* 2015; Stoeck *et al.* 2010), but experience has revealed three major drawbacks in this approach. Firstly, universal eukaryotic primers miss a great part of the diversity and are biased towards certain taxa (Geisen *et al.* 2015a; Jeon *et al.* 2008; Lentendu *et al.* 2014), in particular favouring ciliates and selecting against Amoebozoa (Geisen *et al.* 2015a; Lentendu *et al.* 2014; Fiore-Donno *et al.* 2016). Secondly, a study comparing the outputs of general and cercozoan-specific primers has shown that structuring effects of environmental factors (eg soil fertilization) on communities could be seen only in the datasets generated with specific primers (Lentendu *et al.* 2014). Thirdly, the majority of the reads will stem from multicellular organisms, especially animals and fungi (Baldwin *et al.* 2013; Dupont *et al.* 2016), a waste of sequencing effort when exploring protistan diversity. Thus, using general primers only gives a broad but superficial and biased view of the protistan community composition. For an in-depth understanding of the ecological forces shaping the communities it is advisable to conduct molecular surveys targeting a single monophyletic group. This requires designing and testing primers that will not amplify off-target DNAs while encompassing the whole diversity of the taxon of interest.

It appears from both high-throughput sequencing studies and inventories based on observation that Cercozoa (Cavalier-Smith 1998) often represents, together with Amoebozoa, the dominant protistan group in terrestrial habitats, freshwater and marine ecosystems (Bates *et al.* 2013; Burki & Keeling 2014; Geisen *et al.* 2015b; Grossmann *et al.* 2016; Urich *et al.* 2008). The feeding habits and therefore the ecological roles of such an immense phylum are multiple, including heterotrophs, parasites or autotrophs (Cavalier-Smith & Chao 2003). What makes their study intriguing is the discrepancy between the number of described species (ca. 600) (Pawlowski *et al.* 2012) and the suspected thousands of lineages, a reservoir of hidden diversity that only massive environmental sequencing will reveal (Burki & Keeling 2014). Designing primers for the whole Cercozoa is challenging because of the main dichotomy between Endomyxa (including parasites such as Ascetosporea and Phytomyxea and mostly free-living vampyrellids, *Gromia* and *Filoreta*) and the majority of the taxa grouped under Filosa (Cavalier-Smith & Chao 2003). The monophyly of Endomyxa has not yet been confirmed by phylogenomics: Ascetosporea and *Gromia* + *Filoreta* may be sister to Retaria (Foraminifera, Polycystinea and Acantharea) (He *et al.* 2016; Sierra *et al.* 2016). In Filosa, several main clades can be clearly distinguished by 18S rRNA gene phylogenies, eg Imbricatea, Granofilosea, Thecofilosea, Cercomonadida and Glissomonadida (Bass *et al.* 2009; Howe *et al.* 2011). Although the deep branching of the named ten filosan classes (Bass *et al.* 2009; Cavalier-Smith & Chao 2003) and several new environmental clades are difficult to solve (Bass *et al.* 2009; Cavalier-Smith & Chao 2003; Chantangsi *et al.* 2010; Hoppenrath & Leander 2006; Howe *et al.* 2011), the genetic divergence between the main clades makes the V4 variable region a suitable marker to identify environmental sequences (Bugge Harder *et al.* 2016). Although primers specifically targeting Cercozoa are available they mostly amplify fragments too long to be suitable for MiSeq Illumina (Table 1).

**Table 1.** Primers targeting Cercozoa or cercozoan taxa.

| Specificity | 18S region amplified | Fragment size | Sequencing method | Results | References | MiSeq suitable? |
| --- | --- | --- | --- | --- | --- | --- |
| Cercozoa | nearly complete | 1.26 kb | Sanger | 110 unique sequences | Bass & Cavalier-Smith 2004 | too long |
| Cercozoa | V4 | ca. 270 bp | 454 | 1,585 OTUs 95% | Bugge Harder et al. 2016 | yes |
| Cercozoa | nearly complete | ca. 1kb | Sanger | 24 OTUs 98.7% | Ploch et al. 2016 | too long |
| Cercozoa | V4 and ca. 350 pb upstream | ca. 600 bp | 454 | 187 OTUs, sim. % not stated | Lentendu et al. 2014 | too long |
| *Reticulamoeba* | first third, V4 | 800 bp | Sanger | not stated | Bass et al. 2012 | too long |
| Haplosporidia | V7, V8 & V9 | ca. 670 bp | Sanger | 49 OTUs (> 3 differences) | Hartikainen et al. 2014 | too long |
| Haplosporidia | V7 & V8 | ca. 350 bp | Sanger | 12 "clades" | Pagenkopp Lohan et al. 2016 | yes |
| *Guttulinopsis*,<br>*Helkesimastix* and<br>*Rosculus* | V6 | ca. 480 bp | Sanger | not stated | Bass et al. 2016 | too long |

To test our new primers, we focused on terrestrial environments, following Burki & Keeling (2014): “More than anywhere else, it is in soils that the greatest environmental impact of Cercozoa can be unearthed”. To our knowledge, Cercozoa in biological soil crusts (thereafter, “biocrusts”) have never been specifically targeted to date. Biocrusts play an important ecological role in drylands and extreme habitats, contributing to ca. 50% of the terrestrial biological nitrogen fixation (Elbert *et al.* 2012) and to CO_2_ sequestration, constituting an important reservoir of carbon (Maestre *et al.* 2013). They are formed by living organisms (cyanobacteria, algae, microfungi, lichens and bryophytes) and their by-products, creating a top-soil layer in drylands and as pioneers in disturbed terrestrial ecosystems (Bowker *et al.* 2010).

Our study was aimed at facilitating large-scale, in-depth environmental surveys of Cercozoa by providing newly designed primers suitable for high-throughput sequencing platforms. To ensure reproducible results, we tested our method using DNAs from diverse soils and biocrusts.

## 2. Materials and Methods

### 2.1 Provenance of the soil and biocrust samples and DNA extraction

The provenance of the 26 soil and biocrust samples is detailed in Table S1. Roughly, the samples can be classified in three categories: a) soil samples from mountain grassland collected near Davos (Switzerland); b) biocrusts from Chile (Atacama) and Spitsbergen (Norway) and c) biocrusts from a deciduous mixed forest in Germany. We will refer to them as “Mountain grassland”, “Chile/Spitsbergen” and “Forest”.

Twelve samples were collected for this study, in three ski domains around Davos (Switzerland) in early summer 2015 after the snow melted. The vegetation was removed to access the soil just underneath (5-6 cm depth). Soil samples were kept cold (ca. 4 °C). DNA extraction was conducted within a week time as already described (Fiore-Donno *et al.* 2016).

Within the framework of the German Science Foundation (thereafter DFG) project “Earth Shape”, seven biocrust samples were collected in January 2016 along the Chilean coastline. Sampling sites were chosen to reflect the influence of climate ranging from humid (one sample) to temperate-humid (three samples) and hot desert climate (three samples). Biocrust samples were stored dry until return to Germany, where DNA was extracted by the PowerSoil DNA extraction kit (MoBio Laboratories, Carlsbad, CA, USA) according to the manufacturer’s instructions.

The cold desert sample was collected within the DFG framework “Antarctic Research”, in the project “Biological soil crust algae in the polar regions”, in the Spitsbergen island in Norway. The biocrust was removed from a dark brown soil rich in humic substance, covered with mosses. DNA was extracted following a CTAB protocol using RNase A (Rippin *et al.* 2016) and further purified using the “DNA Clean & Concentrator” (Zymo Research, Irvine, CA, USA).

Within the framework of the DFG Biodiversity Exploratories (Fischer *et al.* 2010), six biocrust samples were collected in separate forest sites in the Schorfheide-Chorin Biosphere Reserve in summer 2014. Biocrusts were only present on litter-free soil at exposed or disturbed sites. Samples were frozen immediately in the field with liquid nitrogen. DNA was extracted from the uppermost three millimeters of the soil crust using the PowerSoil DNA extraction kit (MoBio Laboratories, Carlsbad, CA, USA) according to the manufacturer’s instructions.

### 2.2 Primer design and evaluation *in silico*

To serve as template for primer design, 804 sequences (96% similarity, the longest sequence representative of each cluster) of Cercozoa were downloaded from the PR2 Database (based on GenBank 203 - October 2014) (Guillou *et al.* 2013). After clustering at 96% similarity using vsearch v.1 (Rognes *et al.* 2016) 168 representative sequences were obtained and aligned using MAFFT (Katoh & Standley 2013) with the E-INSi algorithm (gap opening penalty = 3) and refined visually. We designed primers on each side of the variable region V4: two forward and two reverse primers (Table 2). The forward primers have only a base difference between them (S616F_Cerco and S616F_Eocer) at position 660 in our alignment (Appendix S1); S616F_Eocer only match a group of soil environmental sequences related to a genus that could be of importance, *Eocercomonas*, represented by sequence EF023412 (Lesaulnier *et al.* 2008) in our alignment (Appendix S1). Because of too many divergences at the primer sites, our primers could not match the lineages Vampyrellida and Haplosporidia (Appendix S1). To further test the specificity and coverage of our primers and to compare them with existing ones, we used Testprime 1.0 (Klindworth *et al.* 2013) (https://www.arbsilva.de/search/testprime/ last accessed October 2016) on the non-redundant Silva database 128.

**Table 2.**
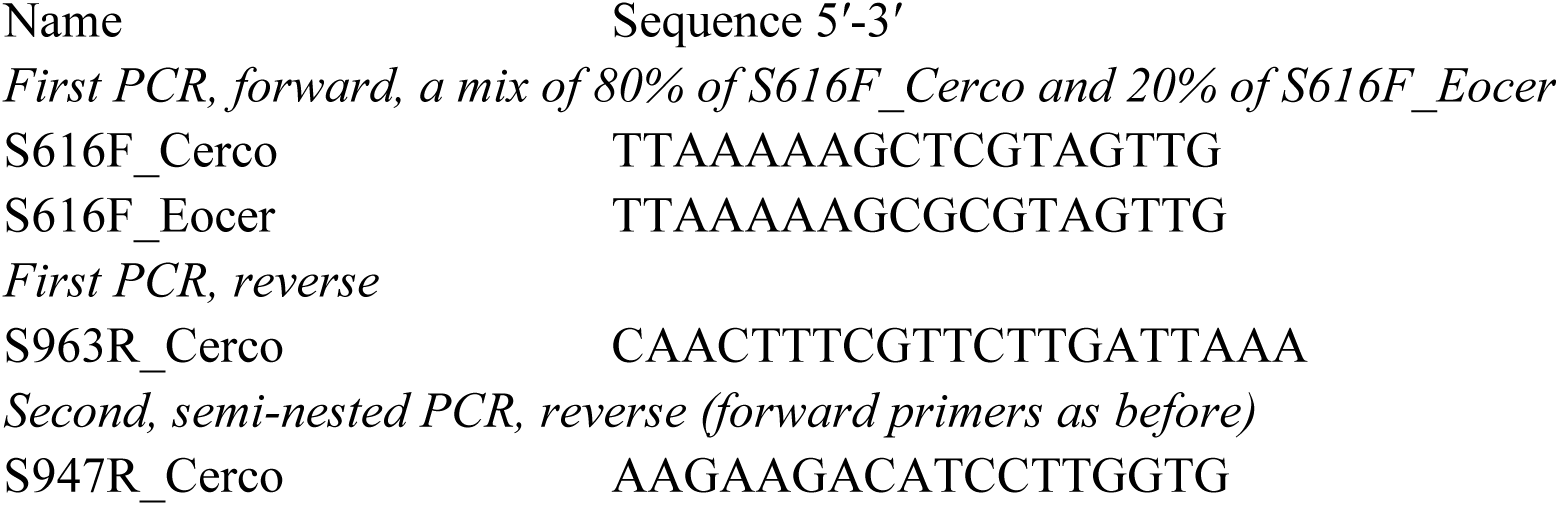
Primers used in this study. The primers used in the second PCR were barcoded (Table S2).

We used a two-step PCR approach with barcoded primers in the second PCR, as recommended to increase reproducibility and avoid diversity biases (Berry *et al.* 2011). The barcodes consisted in eight-nucleotide-long sequences appended to the 5’-ends of both the forward and the reverse primers, because tagging only one primer leads to extensive mistagging (Esling *et al.* 2015). Barcodes were designed to (i) be anti-complementary to the consensus of the reference alignment flanking the primer region; (ii) ensure a balanced nucleotide composition; and (iii) have minimum pairwise edit distances of 3 both for the forward and reverse primers. We designed 16 barcoded versions for the forward S616F_CercoMix and for the reverse S947R_Cerco primers (Table S2), allowing for 256 possible combinations to label samples, although it is advisable to leave at least 10% unused combinations to decrease mistagging (Esling *et al.* 2015). Barcoded primers were specifically ordered for NGS application to Microsynth (Wolfurt, Austria).

### 2.3 PCR amplification, library preparation and sequencing

PCRs were conducted in two steps. In the first PCR the forward primers S616F_Cerco and S616F_Eocer (Table 2) were mixed in the proportions of 80% and 20%, and used with the reverse primer S963R_Cerco. We incorporated 1 μl of 1/10 soil DNA template for the first PCR round and 1 μl of the resulting amplicons as a template for a following semi-nested PCR. We employed the following final concentrations: GreenTaq polymerase (Fermentas, Canada) 0.01 units, buffer 1x, dNTPs 0.2 mM and primers 1μM. The thermal program consisted of an initial denaturation step at 95 °C for 2 min, 24 cycles at 95 °C for 30 s, 50 °C for 30 s, 72 °C for 30 s; and a final elongation step at 72 °C for 5 min.

The number of PCR cycles was kept at 24 since chimera formation arises dramatically after 25 cycles (Michu *et al.* 2010). The second PCR was conducted with barcoded primers, the used combinations are provided in Table S2. All PCRs were conducted twice to reduce the possible artificial dominance of few amplicons by PCR competition (2 × 10 μl for the first and 2 × 27 μl for the second PCR) and the two amplicons were pooled after the second PCR. In parallel, we created an artificial sample of DNAs from known Cercozoa (thereafter, “mock community”), to assist the fine-tuning of the bioinformatic pipeline. The mock community amplicons were obtained from the DNAs of ten cultures with only the second PCR with tagged primers and then purified, quantified with a fluorometer, normalized, pooled and added as a supplementary sample in the Illumina run (Table S3).

The amplicons were checked by electrophoresis and 25 μl of each were purified and normalized using SequalPrep Normalization Plate Kit (Invitrogen GmbH, Karlsruhe, Germany) to obtain a concentration of 1-2 ng/μl per sample. We then pooled the 26 samples and the mock community. During the library preparation amplicons were end-repaired, small fragments were removed, 3’ ends were adenylated, and Illumina adapters and sequencing primers were ligated (TruSeqDNA PCR-Free, Illumina Inc., San Diego, CA, USA). The library was quantified by qPCR, performed following the manufacturer’s instructions (KAPA SYBR^®^ FAST qPCR Kit, Kapabiosystems, Wilmington, MA, USA) on a CFX96 Real Time System (Bio-Rad, Hercules, CA, USA). Sequencing was performed with a MiSeq v2 Reagent kit of 500 cycles on a MiSeq Desktop Sequencer (Illumina Inc., San Diego, CA, USA) at the University of Geneva (Switzerland), department of Genetics and Evolution.

### 2.4 Sequences processing

Paired reads were assembled using mothur v.3.7 (Schloss *et al.* 2009) (which was also used in the following steps) allowing one difference in the primers, no difference in the barcodes, no ambiguities and removing assembled sequences < 100 nt and with an overlap <100 bp (Table 3). Reads were sorted into samples via detection of the barcodes (Table S2). The quality check and removal/cutting of low-quality reads was conducted with the default parameters. Using BLAST+ (Camacho *et al.* 2008) with an e-value of 1-^50^ and keeping only the best hit, sequences were identified using the PR2 database (Guillou *et al.* 2013) and non-cercozoan sequences were removed (Table 3). Chimeras were identified using UCHIME (Edgar *et al.* 2011) as implemented in mothur with a penalty for opening gaps of −5 and a template for aligning OTUs (V4 region of 78 cercozoan taxa, Appendix S2). Sequences were clustered using vsearch v.1 (Rognes *et al.* 2016), with abundance-based greedy clustering (agc) with a similarity threshold of 97% (Table 3).

**Table 3.** Reduction and clustering of reads during the quality filtering.

|  | *Unique* | *Aligned* | *Non Chimeric* | *Genuine Cercozoa* | *OTUs 97% similarity* | *OTUs 97% similarity*<br>*>100 sequences* |
| --- | --- | --- | --- | --- | --- | --- |
| **Environmental reads** |  |  |  |  |  |  |
| Representative sequences | 1,261,799 | 961,591 | 765,248 | 707,796 | 50,905 | 901 |
| All sequences | 2,405,442 | 1,908,247 | 1,600,579 | 1,470,808 | 1,470,808 | 1,266,037 |
| % removed |  | 21 | 16 | 8 | 0 | 14 |
| **Mock community** |  |  |  |  |  |  |
| Representative sequences | 16,402 | 11,246 | 10,707 | 10,626 | 687 | 10 |
| All sequences | 52,318 | 37,904 | 37,094 | 37,009 | 37,009 | 35,563 |
| % removed |  | 28 | 2 | 0.2 |  | 3.9 |

### 2.5 Alignment, evolutionary model and phylogenetic analyses

A critical step in the analyses is aligning the OTUs with as few gaps as possible. Alignments produced without a template resulted in more than 10,000 positions because of the length variation of the V4 region (320-345 bp). A more reliable alignment could be obtained using mothur with the template for the V4 region provided here (Appendix S2). Eleven OTUs had a low alignment score and were discarded. The aligned OTUs were added to our reference alignment augmented with sequences identified as the best Blast hit for each OTU (in total 178 reference sequences included in Appendix S1), using mothur as described above. We conducted phylogenetic analyses first on the assemblage of the 178 reference sequences, to check for its suitability for identifying OTUs. The alignment had 1,386 positions, 946 alignment patterns, and a proportion of undetermined characters of 9.22%. The alignment with the 890 OTUs partial sequences added resulted in 1,059 sequences, 1,386 positions, 994 alignment patterns, and a proportion of undetermined characters of 71.71% (Appendix S1). Maximum likelihood (ML) analyses were run using RAxML 8.2.4 (Stamatakis 2014), with the GTR model of substitution and a 25 rate category discrete gamma distribution. The best scoring ML tree was inferred from 400 randomized starting Maximum Parsimony trees using the GTRMIX model. The best-scoring tree was used to report the confidence values as percentages obtained through 400 non-parametric bootstraps. The tree was saved using iTOL v3.4 (Letunic & Bork 2016).

### 2.6 Statistical analyses

To evaluate if more sequencing effort would have revealed more richness, we carried out a rarefaction analysis with the vegan package, function specaccum (Oksanen *et al.* 2013) in R v.3.1.2 (R Development Core Team 2014) (Fig. S1). To estimate intra- and inter-site category similarities, Venn diagrams were calculated with the VennCounts and VennDiagram function in the Limma library (Bioconductor Project) (Smyth 2005); the multiple-site Simpson-based similarity indices were obtained using the R script provided in Baselga *et al.* (2007), with presence-absence tables as input.

## 3. Results

### 3.1 Sequencing and bioinformatic pipeline

During the sequencing the cluster density was 904 K/mm^2^ and the overall quality high (95.22% ≥ Q30). A percentage of reads with unused barcode combinations were present as a consequence of mistagging during the Illumina sequencing run, amounting to 2.72%. The mock community was analysed first. We could retrieve the expected ten OTUs at 97% similarity when deleting OTUs represented by <100 sequences, thus we used these settings for the main analysis. Without this cutoff we would have obtained 687 OTUs from an assemblage of 10 amplicons (Table 3), proving that the error rate is certainly more detrimental than usually considered and might conduct to an overestimation of the diversity. We advocate here for including whenever possible a mock community in every run. From the soil/biocrust samples we obtained 901 OTUs, representing 1,266,037 sequences (Table 3 and Appendix S1). The length of the amplified V4 region (without primers) varied from 320 to 345 bp. A database with the abundance of each OTU per site and their taxonomic assignment is provided (Table S4).

### 3.2 Primers evaluation *in silico*

*In silico* PCR performed on the non-redundant Silva database showed that our primers combination would not amplify any prokaryote, nor plants or animals, and only one fungal sequence. In the SAR clade, except one ciliate and four diatoms, all recovered sequences belonged to Cercozoa. 68% of the cercozoan sequences were matched *in silico* (Fig. 1).

**Figure 1.**
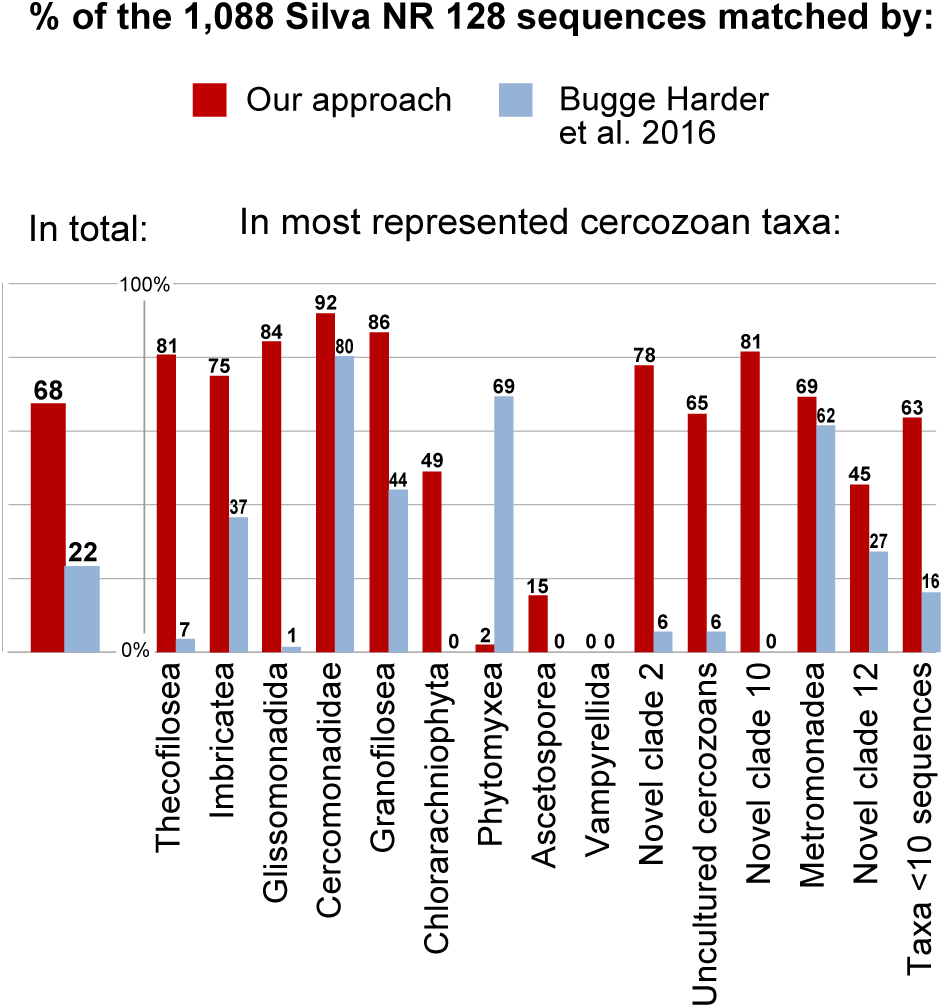
Comparaison of the *in silico* amplification efficiency of our new primers and those of Bugge Harder *et al.* (2016), on the 1,088 cercozoan sequences from the Silva nonredundant database. Only high-rank tanomic names are shown. Taxa represented by less than 10 sequences are grouped.

Comparison of our primers with Cerc479F and Cerc750R (Bugge Harder *et al.* 2016) (the only MiSeq compatible Cercozoa-specific primers, Table 1) revealed that the latter could match *in silico* only 248 cercozoan sequences, missing 78% of the database sequences, overall missing Chlorarachniophyta, Novel Clade 10 and Ascetosporea (Fig. 1). Our primers performed better *in silico* on very abundant and widespread taxa like Thecofilosea and Glissomonadida, and were only outcompeted when targeting the parasitic Phytomyxea. Both approaches missed the parasitic and very divergent Vampyrellida and Haplosporidia (the latter in Ascetosporea) (Fig. 1 and Appendix S1).

### 3.3 Cercozoan alpha diversity

The 901 OTUs represented 212 unique Blast best hits. Only 43% of the OTUs were 97-100% similar to any known sequence (Fig. S2). At a high taxonomic level, the majority of the OTUs could be assigned to Filosa (96%) and only 2% to Endomyxa, the remaining reads were assigned to “undetermined Cercozoa or Filosa” (that is, environmental sequences that could not be assigned more precisely) and the *incertae sedis* Novel Clade 12 (Fig. 2A). In Filosa, the majority of the OTUs could be assigned to the classes Sarcomonadea (Glissomonadida and Cercomonadida), Imbricatea and Thecofilosea. At the level of orders, the highest diversity was retrieved in Glissomonadida followed by Euglyphida (Imbricatea), Cryomonadida (Thecofilosea) and Cercomonadida (Fig. 2B). The 23 most abundant OTUs (> 10,000 sequences) accounted for 54% of the total sequences, while many small OTUs (758 < 1,000 sequences) contributed only to 17% of all sequences. The most abundant OTU were assigned to the *Rhogostoma*-lineage, the family or genus Sandonidae and *Sandona*, the species *Corythion dubium*, *Assulina muscorum*, *Neoheteromita globosa* and *Flectomonas ekelundi*, three species of *Euglypha* and undetermined Trinematidae, Cercomonadidae and Allapsidae (Table S4). The rarefaction curve reached a clear plateau, suggesting that < 100′000 sequences would have been sufficient to reflect the global OTUs richness (as revealed by our primers) at all sites (Fig. S1). This was also true, to a lesser extent, when the three sites categories were considered individually (Fig. S1). The phylogenetic tree obtained with the 178 reference sequences was mostly consistent with published phylogenies (eg Howe *et al.* 2011), allowing to place OTUs in a phylogenetic scheme with reasonable confidence Fig. S3). The tree was rooted with Proteomyxidea and Phytomyxea (Endomyxa) (46% bootstrap support), the Novel Clades 10-11-12 were paraphyletic to a monophyletic Filosa (86%). In Filosa, the following main clades were recovered: Chlorarachnea (100%), Granofilosea (28%), Metromonadea (70%), Paracercomonadidae (73%), Cercomonadidae (70%), a clade of *incertae sedis* “uncultured Filosa” (86%), Glissomonadida (including Pansomonadidae, 58%), Thecofilosea (including Phaeodarea, 14%) and Imbricatea (7%). In the second phylogenetic tree (Fig. S4) our new environmental sequences were present in nearly every clade of the tree, including the basal clades Phytomyxea and Novel Clades 1011-12, but were absent from Chlorarachnea and Phaeodarea (both marine) and Vampyrellida (the latter not matched by our primers).

**Figure 2.**
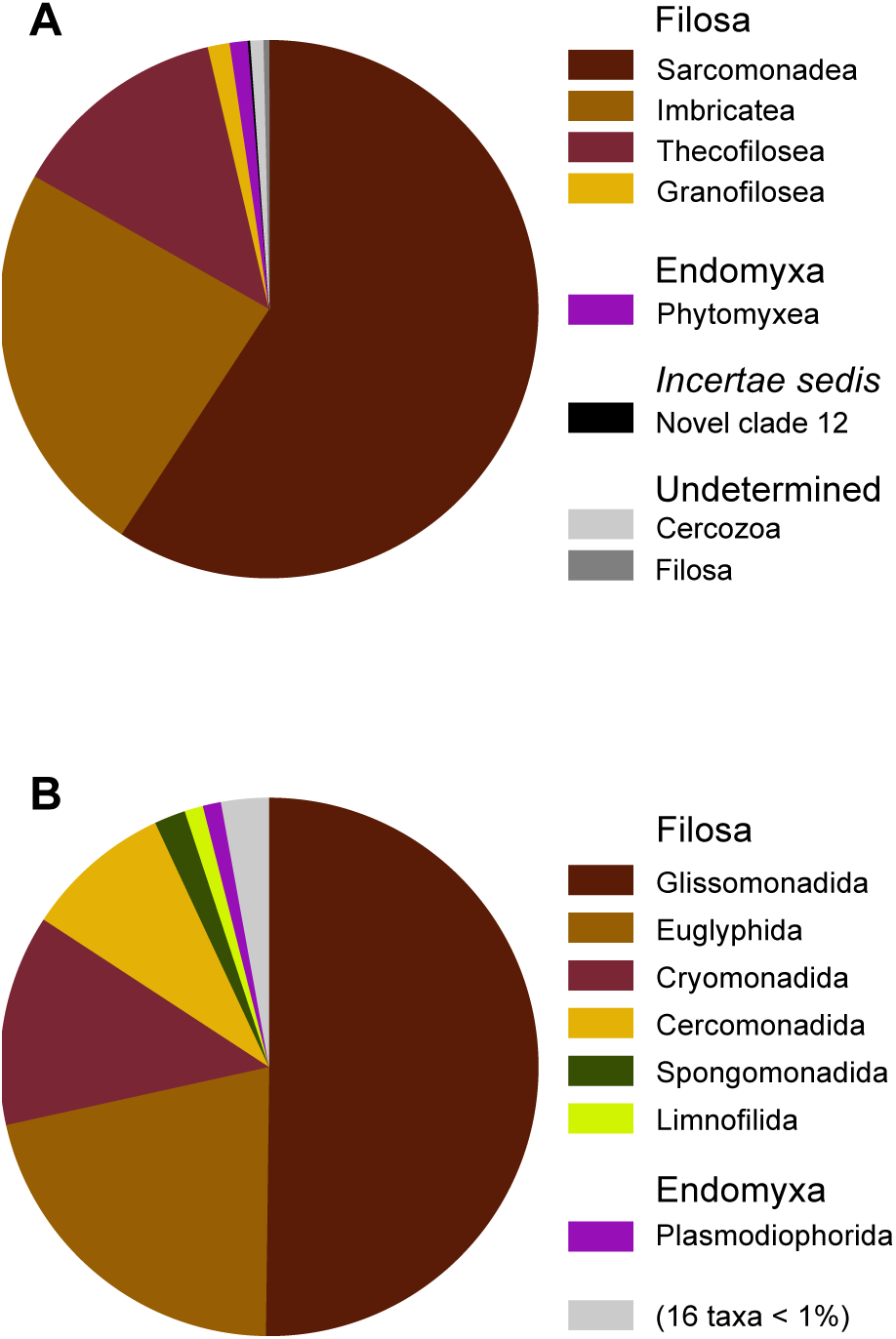
Relative contribution of the OTUs to the taxonomic diversity. Taxonomical assignment is based on the best hit by BLAST. **(A)** At the level of the classes. **(B)** At the level of the orders. “Undetermined” refer to environmental sequences in the reference database that could not be assigned to the next lower-ranking taxon.

In the four most abundant classes, ie Sarcomonadea (Glissmonadida and Cercomonadida), Imbricatea, Thecofilosea and Granofilosea (Fig. 2), the dominant genera were *Euglypha* (69 OTUs) and *Cercomonas* (60 OTUs); the vast majority of reads corresponded to sequences from environmental samples that themselves did not match any known sequence in the database (Table S5). The highest percentage of such undetermined lineages was found in Thecofilosea (91%), followed by Granofilosea (73%) and Glissomonadida (70%). It was in Glissomonadida that the highest number of sequences that could be assigned at the family level only was found: 144 Sandonidae, 94 Allapsidae, while only 12 could be assigned at the order level (Table S5).

We found an OTU (representing 113 sequences) related to Marimonadida (Imbricatea), an order described as marine (Howe *et al.* 2011) and six OTUs related to *Paulinella* (89.9 - 94.6% similarity by Blast), a freshwater genus (Meisterfeld 2000) (Table S4). Our six *Paulinella*-related OTUs were extremely abundant in two biocrust samples S01 (temperate forest, > 6,000 sequences) and Spitsbergen (1,732 sequences). We found many OTUs related to *Limnofila*, a mostly freshwater genus (Bass *et al.* 2009) and many *Metopion*-related OTUs (Table S4). *Metopion* is a predator on other eukaryotes and as such not easily detected in culture surveys (Howe *et al.* 2011), thus it is important to detect its presence by ePCR .

We found eight OTUs for a total of 2,371sequences (Table S4) related to two sequences labelled as “Filosa” in the PR2 database - EF024287 (Cercomonadida environmental sample clone Elev_18S_684) and AB534506 (Uncultured eukaryote, clone: I_3_96), branching at the base of Glissomonadida with low support (Fig. S3 & S4) and having 95% identity to each other. When searching GenBank for close relatives (https://blast.ncbi.nlm.nih.gov/Blast.cgi?PROGRAM=blastn&PAGE_TYPE=BlastSearch&LINK_LOC=blasthome,blastn_2.6.1+,_nt/nr_database, last accessed May 2017), we could not find matches with >93% identity, and the ten best were of environmental origin.

### 3.4 Cercozoan beta diversity

All 26 DNA samples gave positive results. The average number of sequences per site was 48,693 (maximum: 121,343, minimum 4,648) and the average number of OTUs per site was 332 (maximum 442, minimum 123). Our sequencing depth was sufficient, as reflected by the levelling of the rarefaction curves (Fig. S1) and the high number of OTUs per sample (Table S4). Although the number of OTUs was similar between the three site categories, from 732 to 768, their taxonomic repartition was uneven, even for the most abundant orders. For example, Cercomonadida was somewhat relatively more abundant in the mountain grasslands and Euglyphida in the Chile/Spitsbergen biocrusts (Fig. 3). The less frequent orders showed more marked preferences, with Tectofilosida more abundant in the forest biocrusts, Novel Clade 12 in the mountain grasslands and “undetermined Marimonadida” found only in two humid temperate *Araucaria* forest sites (Na_H and Na_V, Table S1 & Fig. 3). Nonetheless, the number of shared OTUs between sites was high (61%), while the proportion of OTUs unique to each site category was low (4 - 6%) (Fig. 4). The Simpson-based multiple-site similarity indices showed that the communities were more similar inside each site category than between sites, the lowest index being associated with the three sites taken together (Fig. 4).

**Figure 3.**
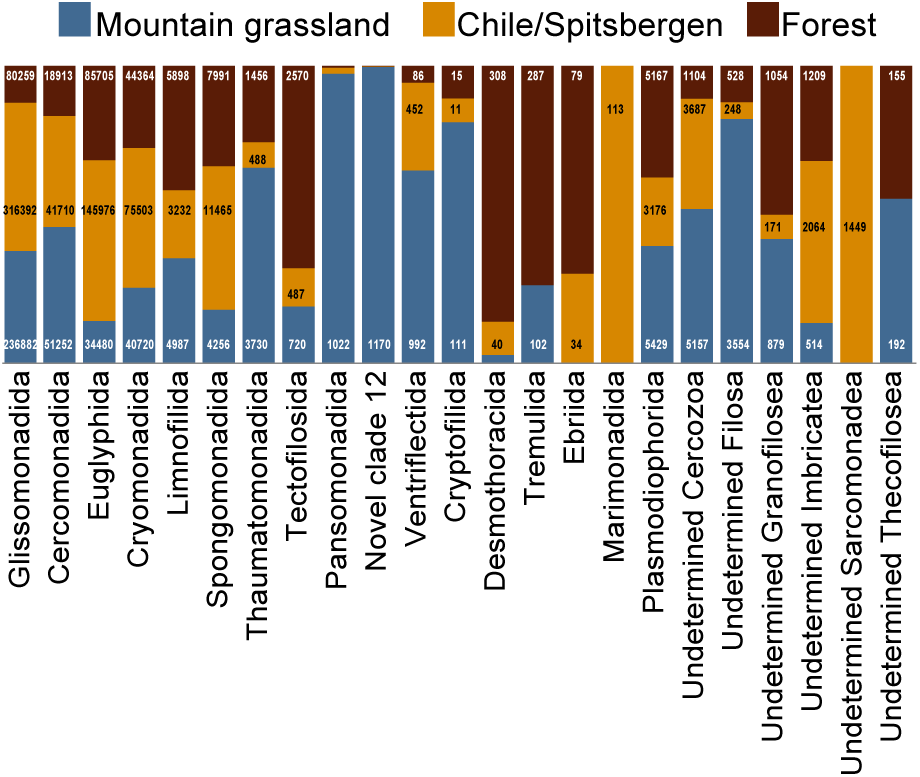
Proportions of OTUs (grouped by orders) in each site category, with the corresponding number of sequences. The distribution is not even, suggesting biogeographic patterns.

**Figure 4.**
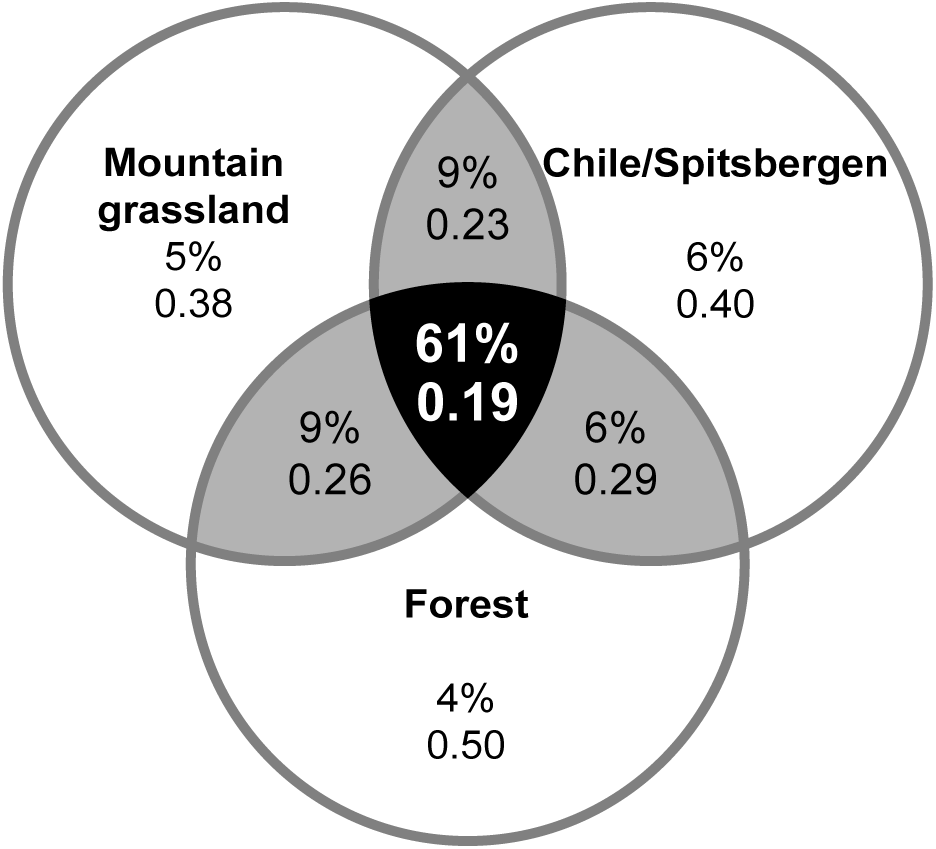
Inter-site categories similarities: Percentages of unique OTUs per region (in the circles) and shared between regions (in the intersections). Shaded in black, OTUs shared by the three categories (as percentage of the sum of the three regions); shaded in grey, OTUs shared between two regions (percentage of the sum of the two regions). Simpson-based multiple-site similarity indices, calculated for each and between site categories are given below the percentages.

## 4. Discussion

Our newly designed primers fulfilled the requirements of being highly specific (only 8% of non-cercozoans sequences, Table 3), while the 901 retrieved OTUs (97% similarity) spanned across the diversity of Cercozoa (Fig. S4). Their main limitation consisted in missing most of the divergent lineages that could be more related to Retaria than Cercozoa: Ascetosporea, Gromiidea and Reticulosida (He *et al.* 2016; Sierra *et al.* 2016). We could not retrieve any of those taxa from the environment in spite of the *in silico* test showing some matches for *Gromia* and for 15% of the Ascetosporea. The low proportion of OTUs assigned to Phytomyxea (2%, 11 OTUs in Plasmodiophorida, 13,772 sequences) (Table S4) probably underestimated their actual diversity, since our primers matched only 2% of the phytomyxean sequences *in silico* (Fig. 1). In Filosa, our primers could not detect from soil/biocrusts samples the marine Phaeodaria (Thecofilosea) (Takahashi & Anderson 2000) and Chlorarachnea, although they are matched by our primers, that could thus also be used in marine environments (Fig. S4).

The use of barcoded primers, in addition of being an interesting economical alternative to multiple libraries, allowed to detect mistagging happening during the Illumina sequencing run: in this case, we could estimate it to a 2.72% of the reads. The mixing of adjacent sequencing cluster of reads can happen when the density in the flow cell is higher than recommended, that is >865-965 K/mm^2^ (https://www.illumina.com/systems/sequencing-platforms/miseq/specifications.html, last accessed April 2017) - which was not the case here (904 K/mm^2^). The mistagging issue is a potential source of confusion in the ecological evaluation of the beta diversity, and its influence should be assessed for every run.

Our primers are suitable for in-depth cercozoan diversity surveys, since they revealed a still unearthed diversity, with 57% of the OTUs not assignable to a known sequence at >97% similarity. Concomitantly, this shows that efforts to increase the reference database are needed. The dominating taxon, Glissomonadida, also had the highest proportion of OTUs phylogenetically distinct from characterized lineages (Table S5 and Fig. S4). It is generally agreed that small flagellates belonging to Glissomonadida (predominantly gliding bacterivores - except Viridiraptoridae, amoebae and parasites) are the most abundant in various kinds of soils worldwide (Bates *et al.* 2013) (therein incorrectly referred to as Heteromitidae), in temperate grasslands and forests (Domonell *et al.* 2013; Howe *et al.* 2009) and in heathlands (Bugge Harder *et al.* 2016). This astounding diversity sparks off questions about the role they play in the microbial food web and in biogeochemical soil processes.

The second most abundant taxon, Euglyphida, comprises species that are typically abundant in peat bogs, and generally present in soil and freshwater habitats (Geisen *et al.* 2015b; Lara *et al.* 2007). Most OTUs assigned to the family Trinematidae in our tree display long branches, suggesting a high potential of still unknown taxa. *Euglypha* might be a significant player in the soil silica cycle, as important as plants, in an interaction involving mycorrhizae in the rhizosphere of ericoids (Wilkinson 2008).

It was more surprising to find so many Cryomonadida (in the class Thecofilosea - mostly thecate amoebae but also flagellates) (Dumack *et al.* 2017), since they were not mentioned in most soil microscopical inventories (Domonell *et al.* 2013; Esteban *et al.* 2006) and were not considered abundant in soil by Bugge Harder *et al.* (2016) nor by a transcriptomic study of various soils (Geisen *et al.* 2015b). Environmental sequences were retrieved firstly from marine habitats (Bass & Cavalier-Smith 2004), a few were later found in soil and faeces (Bass *et al.* 2016). Our data clearly showed the existence of an unprecedented diversity of the *Rhogostoma*-lineage in soils and biocrusts of various geographic regions. Interestingly, it has been found that *Rhogostoma* populations sharply increased when a carbon source (glycerol) was added to a membrane bioreactor treating landfill leachates (to remove nitrates), so that *Rhogostoma* may become dominant in fields enriched in available carbon and nitrogen (Remmas *et al.* 2017). In a fish farm, a massive outbreak of the nodular gill disease of the rainbow trout was observed, associated with a high abundance of *Rhogostoma minus* in the gills of affected fish (Dyková & Tyml 2016). These new findings, along with our discovery of the abundance of *Rhogostoma*-lineage species in soil and biocrusts, suggests that members of this genus may play a prominent and diversified role in nutrient cycling (Dumack *et al.* 2017), as well as being opportunistic pathogens.

Interestingly, we could detect OTUs related to lineages considered as mainly freshwater (*Limnofila*, *Paulinella*) or even marine (Marimonadida) (Table S4): perhaps, for a protist there is not much difference between dwelling in water trapped between soil particles or in puddles, pools, ponds or lakes. Our primers could also detect a potential new lineage, a clade labelled in our tree as “undetermined Filosa”, that could not be related to any known taxa and with no discernible phylogenetic affiliation.

Next generation sequence surveys of eukaryotes have revealed that Cercozoa are extremely diverse and abundant, with a large and yet undiscovered component, both in marine and terrestrial ecosystems. Our study revealed widely distributed and highly diverse cercozoan communities in soils and biocrusts in very diverse climatic regions. Our primers and protocols will now allow large-scale, multi-sample cercozoan environmental diversity studies, due to the in-depth scanning power of the here described specific barcoded primers compatible with multiplex MiSeq Illumina sequencing. This will enhance our fundamental understanding on organismal and genetic diversity, on the drivers structuring cercozoan communities and particularly on their functional role in all types of ecosystems.

## 5. Acknowledgments

We thank Franck Lejzerowicz for designing the barcodes for the primers and helpful suggestions; Laura Mehner for her contribution in the primers and testing design; Sebastian Flues, Kenneth Dumack and Sebastian Hess for providing the DNAs of the mock community; Peter Leinweber for collecting samples in Chile. This work has been funded by the DFG Priority Program 1374 “Infrastructure-Biodiversity-Exploratories” (BO 1907/2-2). E. Samolov, K. Glaser and U. Karsten appreciate funding by the DFG projects Crustweathering (KA899/32-1, Priority Program “EarthShape”) and Crustfunction I (KA899/28-1, Priority Program 1374 Infrastructure-Biodiversity-Exploratories). At the University of Geneva, we thank Jan Pawlowski, Emanuela Reo and Ewan Smith and we are liable to the Swiss National Science Foundation Grant 316030 150817 funding the MiSeq instrument. We thank Markus Fischer, Eduard Linsenmair, Dominik Hessenmöller, Jens Nieschulze, Daniel Prati, Ingo Schöning, François Buscot, Ernst-Detlef Schulze, Wolfgang W. Weisser and the late Elisabeth Kalko for their role in setting up the Biodiversity Exploratories project. Field work permits were issued by the responsible state environmental office of Brandenburg (according to § 72 Brandenburgisches Naturschutzgesetz). The funders had no role in the study design, data collection and analysis, decision to publish, or the preparation of the manuscript.

## 7. Data accessibility

Raw sequences have been deposited in GenBank, BioProject PRJNA360862 and the 901 OTUs (represetative sequences) under accession numbers KY553293-KY554193.

## 8. Supporting Information

Additional Supporting Information may be found in the online version of this article:

**Appendix S1**. Alignment of cercozoan nearly complete small subunit ribosomal RNA gene sequences, in fasta format. The alignment includes 178 reference sequences with GenBank accession number and taxonomic assignment and our 901 OTUs (V4 partial sequences). First sequence: mask showing the unambiguously 1,386 aligned positions. Second to fourth sequences: primers designed in our study and designed by Bugge Harder *et al.* (2016), targeting the V4 region. Fifth to eighth sequences: consensus at 96 and 98% similarity, with and without Vampyrellidae.

**Appendix S2**. Reference alignment of the V4 region used as template for aligning the OTUs (78 sequences, fasta format).

**Figure S1**. Rarefaction curves describing the observed number of OTUs as a function of the sequencing effort, all sites pooled and the three site categories separately. Saturation, when all sites were pooled, was reached with < 100,000 sequences.

**Figure S2**. Similarities of the OTUs with known sequences. OTUs are classified according to their percentage of similarity to the next kin by BLAST. The horizontal bar length is proportional to the number of OTUs in each rank.

**Figure S3**. Cercozoa Maximum Likelihood phylogenetic tree of the 178 reference sequences. Alignment = 1,386 positions. The tree is rooted between Endomyxa and the remaining taxa. Main clades are named according to the PR2 database taxonomy.

**Figure S4**. Cercozoa Maximum Likelihood phylogenetic tree. Alignment = 1,068 taxa, 1,386 positions. The tree is rooted between Endomyxa and the remaining taxa. Labels colors: Red = OTUs found in this study; Black = the 178 reference sequences; Gray = clades for which no OTUs were found. Main clades are named according to the PR2 database taxonomy.

**Table S1**. Place of collection and charateristics of the samples (pdf).

**Table S2**. **A.** Primers and barcodes designed for multi-sample Illumina runs targeting Cercozoa, allowing for 256 combinations. Consensus sequences: see Appendix S1. **B.** Combinations of barcodes used in this study, with the corresponding samples (pdf).

**Table S3**. Species name, taxonomic assignment, GenBank accession number of the closest match of the sequences obtained and providers of the ten cercozoan cultures composing the mock community (pdf).

**Table S4**. Database of the distribution and abundance of each OTU per site, followed by the taxonomic assignment, the GenBank accession number of the best hit by BLAST and the percentage of similarity (PR2 database) (excel).

**Table S5**. Taxonomic assignment (PR2 database) at the genus level. The four most abundant classes (Sarcomonadea divided into its two major orders) are represented. Number of OTUs per taxon and percentages of “Undetermined” taxa (environmental sequences) are given. The latter percentages indicate that the class/order is understudied or undersampled and that our study revealed a hidden diversity (pdf).

## 9. Author Contributions

AMFD and MB conceived the study, AMFD designed the primers, performed the work and wrote the article; MR, BB, ES, KG, UK provided DNAs; CR collected and provided soil samples; all authors edited the manuscript.

